# Experience alterations in white matter structure and myelin-related gene expression in adult rats

**DOI:** 10.1101/532572

**Authors:** Cassandra Sampaio-Baptista, Astrid Vallès, Alexandre A. Khrapitchev, Guus Akkermans, Anderson M. Winkler, Sean Foxley, Nicola R. Sibson, Mark Roberts, Karla Miller, Mathew E. Diamond, Gerard J.M. Martens, Peter De Weerd, Heidi Johansen-Berg

**Affiliations:** Wellcome Centre For Integrative Neuroimaging, FMRIB, Nuffield Department of Clinical Neurosciences, University of Oxford, UK; Department of Molecular Animal Physiology, Donders Institute for Brain, Cognition and Behaviour, Radboud Institute for Molecular Life Sciences (RIMLS), Radboud University Nijmegen, 6525 GA Nijmegen, The Netherlands; Department of Neurocognition, Faculty of Psychology and Neurosciences, Maastricht University, 6200 MD Maastricht, The Netherlands; Cancer Research UK and Medical Research Council Oxford Institute for Radiation Oncology, Department of Oncology, University of Oxford, Churchill Hospital, Oxford OX3 7LE, UK; Department of Cognitive Neuroscience, Radboud University Nijmegen, Donders Institute for Brain, Cognition and Behaviour, 6500 HB Nijmegen, The Netherlands; Maastricht Centre for Systems Biology (MaCSBio), Maastricht University, Maastricht, The Netherlands; Tactile Perception and Learning Lab, International School for Advanced Studies (SISSA), Trieste, Italy

## Abstract

White matter (WM) plasticity during adulthood is a recently described phenomenon by which experience can shape brain structure. It has been observed in humans using diffusion tensor imaging (DTI). However, it remains unclear which mechanisms drive or underlie WM plasticity in adulthood. Here, we combined DTI and mRNA expression analysis and examined the effects of somatosensory experience in adult rats. Somatosensory experience resulted in differences in WM and grey matter structure. C-FOS mRNA expression, a marker of cortical activity, in the barrel cortex correlated with the structural WM metrics, indicating that molecular correlates of cortical activity relate to macroscale measures of WM structural plasticity. Analysis of myelin-related genes revealed higher myelin basic protein expression in WM, while genome-wide RNA sequencing analysis identified 134 differentially-expressed genes regulating proteins involved in functions related to oligodendrocyte precursors proliferation and differentiation, regulation of myelination and neuronal activity modulation. In conclusion, the macroscale measures of WM differences identified in response to somatosensory experience are supported by molecular evidence, which strongly suggest myelination as, at least, one of the underlying mechanisms.

**SIGNIFICANCE STATEMENT:** White matter plasticity is a recently described mechanism by which experience shapes brain structure and function during adulthood. This phenomenon was first described in adult humans with complex motor skill learning using whole brain non-invasive diffusion tensor imaging (DTI). Here we report structural changes in white matter detected with DTI after novel somatosensory experience in rats. We further support these findings with mRNA evidence of differentially expressed genes involved in functions compatible with regulation of cell proliferation, myelin thickness and neuronal modulation. These putative molecular mechanisms offer insights about the underlying DTI correlates of experience-dependent WM plasticity and provide further evidence for myelin plasticity in adulthood.

## INTRODUCTION

There is accumulating evidence that structural changes in white matter (WM) occur in response to changes in experience, even during adulthood (Sampaio-Baptista and Johansen-Berg, 2017). For example, neuroimaging studies have reported experience-induced structural WM plasticity in both humans (Scholz et al., 2009; Hofstetter et al., 2013) and rodents (Blumenfeld-Katzir et al., 2011; Sampaio-Baptista et al., 2013). However, it is unclear how such macroscale changes relate to the underlying molecular mechanisms of experience-dependent WM plasticity during adulthood.

Structural changes in barrel cortex, such as synapse and spine formation (Trachtenberg et al., 2002) have been extensively reported in response to whisker stimulation (Knott et al., 2002), deprivation (Holtmaat et al., 2006) and somatosensory learning (Kuhlman et al., 2014), along with corresponding changes in expression levels of genes, such as BDNF (Rocamora et al., 1996) and synaptophysin (Ishibashi, 2002). More recently, somatosensory enrichment in middle-aged rats resulted in an increase in oligodendrocyte numbers and integration in barrel cortex (Hughes et al., 2018). However, it is not clear whether somatosensory experience also induces WM plasticity or whether such structural plasticity is localized only to cortical regions.

Here, we investigated if somatosensory experience in adult rats induces WM plasticity at the macro and the molecular scale, by combining neuroimaging and mRNA expression analysis. Adult rats were trained in a texture detection task (TDT) (von Heimendahl et al., 2007) and WM structure was assessed with diffusion tensor imaging (DTI). Additionally, we analysed mRNA expression to further support the structural findings and identify putative molecular mechanisms underlying experience-dependent WM plasticity.

## RESULTS

Three months old rats (n=28) were trained on a texture detection task (TDT) (von Heimendahl et al., 2007) that required a texture identity (rough or smooth) to be associated with a reward side (e.g., turn left on rough, right on smooth). After the initial shaping period, it took the trained animals between 5 and 17 days to reach criterion performance of 2 sessions > 80% accuracy (Fig.1A, B).

**Fig. 1.**
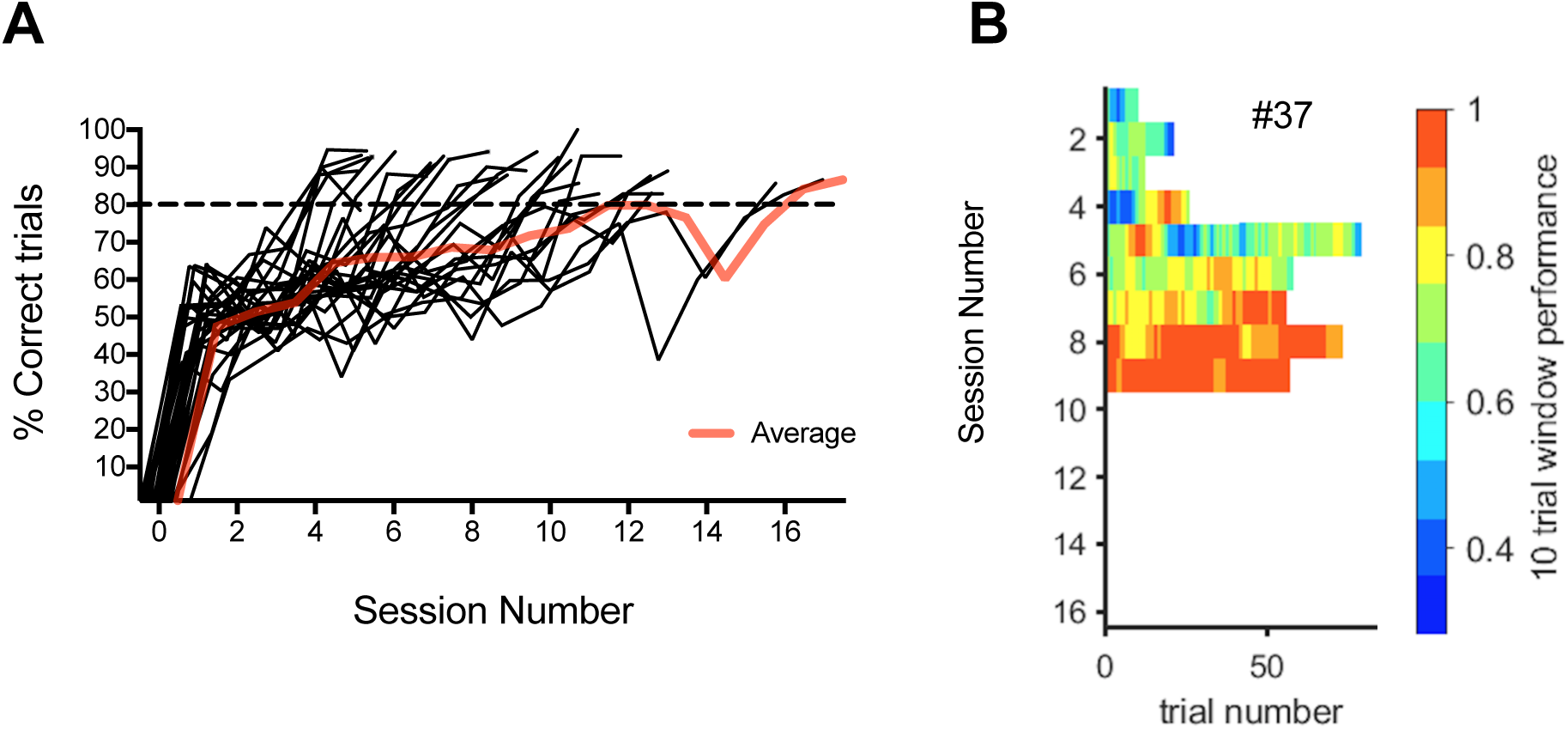
Texture Detection Task (TDT) Performance. **A** Individual performance accuracy (% correct trials) of animals in the TDT group trained on the P100 texture (n=28). Red line represents average group accuracy. **B** Running average graph of an example rat displaying accuracy per day, with each day represented by a horizontal bar showing color-coded performance levels over 10 trial windows. The total number of trials per session (∼30 minutes session) tends to increase and performance improves over days. Colour-coding performance scale shown on the right.

## NEUROIMAGING ANALYSIS RESULTS

### DTI multimodal analysis shows WM structural differences between groups and correlations with performance in the texture detection task

To assess effects of experience on WM microstructure we jointly analyzed structural measures calculated from post-mortem DTI scans of left hemispheres from TDT (n=28) and passive control (PC, n=20) animals. The PC group was handled daily for a few minutes, without any exposure to the testing setup. We performed a non-parametric combination (NPC) for joint inference analysis, as implemented in PALM, over the 4 DTI measures (fractional anisotropy (FA), mean diffusivity (MD), radial diffusivity (RD) and axial diffusivity (AD)), using the Fisher’s combining function. We tested for a concordant direction of effects across all measures, while allowing the assessment of the significance maps for each measure separately (reported in the partial tests), and as such, inference on which measures would drive a significant joint result, with correction for the multiplicity of tests (Winkler et al., 2016) (see Methods for more details).

We first tested for between-group differences across all voxels in the WM skeleton. A significant cluster was found with the NPC joint inference analysis (p < 0.05, fully corrected across all voxels, Fig. 2A) covering a large area of WM under prefrontal and sensorimotor regions (Fig. 2A). These are relevant WM regions for the texture detection task, in the light of the barrel cortex’s role in somatosensory information processing (for review see (Feldmeyer et al., 2013)).

**Fig. 2.**
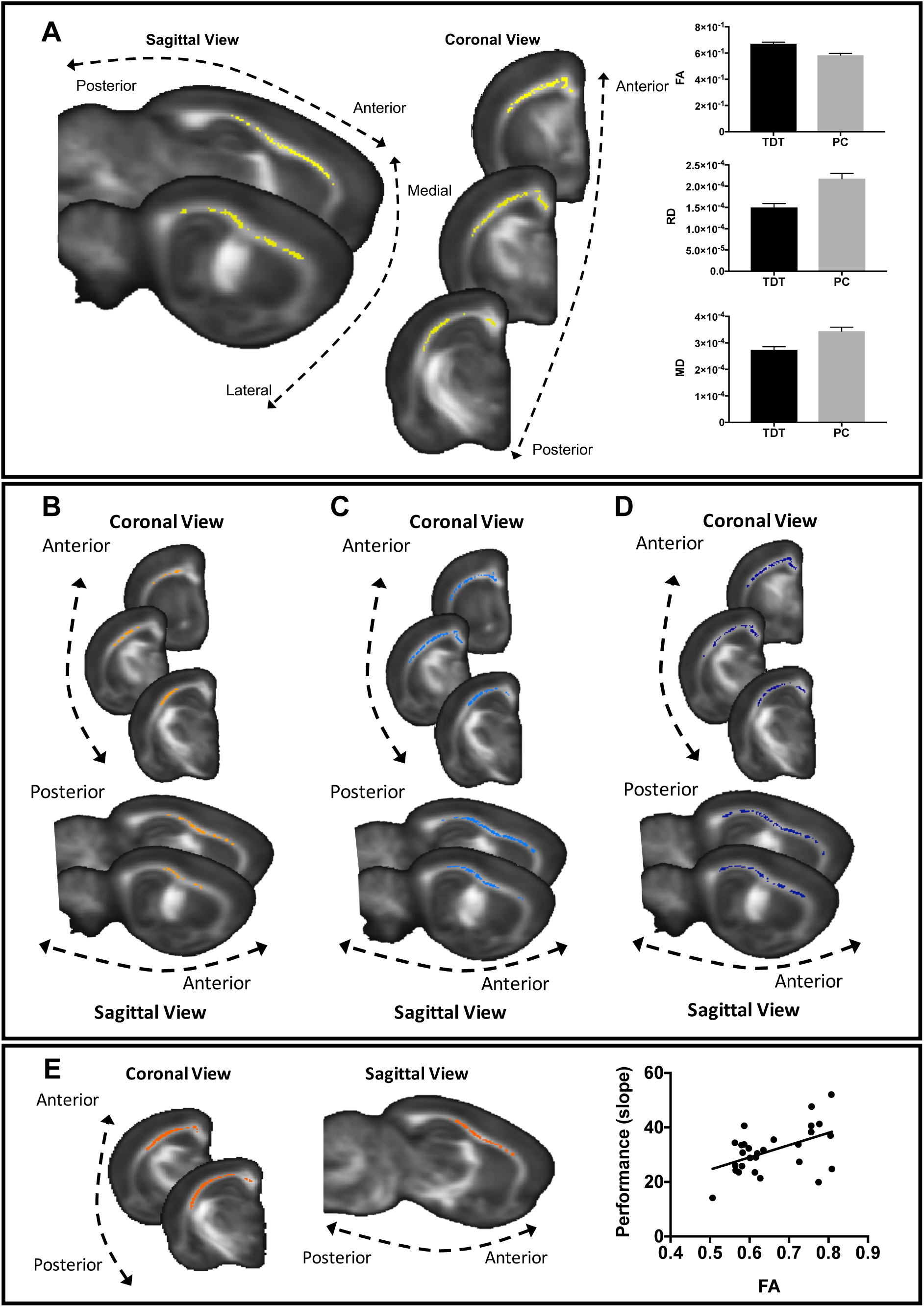
DTI analysis of WM structure. **A** NPC Fisher’s test for joint inference over the 4 DTI measures (FA, MD, RD and AD) (cluster in yellow) was found to be significant p < 0.05 (fully corrected). Bar graphs of FA, RD and MD estimated marginal means (adjusted for the number of exposure days used as a covariate in the model) of the significant yellow cluster are shown to illustrate the direction of differences and not for inference. Error bars represent standard error. **B-D NPC partial tests. B** FA (in orange) was significantly higher in the TDT group (p < 0.05, fully corrected). **C** MD (in blue) was significantly lower in the TDT group (p < 0.05, fully corrected). **D** RD (in blue) was significantly lower in the TDT group (p < 0.05, fully corrected). **E** Performance rate correlates with FA (cluster in red) (p < 0.05, fully corrected). Scatter plot showing the correlation between mean values of the significant clusters and performance rate is displayed for visualisation of the range of values only and not for inference. In all panels, significant clusters are superimposed on the mean FA template.

Additionally, The NPC partial tests showed that, compared to controls, the TDT group had significantly higher FA (p < 0.05, fully corrected across all voxels and the 4 measures) (Fig. 2B), lower MD (p < 0.05, fully corrected across all voxels and the 4 measures) (Fig. 2C), and lower RD (p < 0.05, fully corrected across all voxels and the 4 measures) (Fig. 2D) across similar areas of WM. AD was not significantly different between groups (p = 0.842, fully corrected across all voxels and the 4 measures).

Secondly, to assess the relationship between task performance and the neuroimaging structural measures, we again used a NPC Fisher’s joint inference analysis to test for voxel-wise correlations between 4 DTI measures and performance rate (slope of the individual curves) across individual animals (n = 28). We did not find a significant correlation between performance rate and the joint 4 DTI measures. However, there was a significant correlation between performance rate and FA (p < 0.05, fully corrected across all voxels and the 4 measures, Fig. 2E) and a trend for RD (p = 0.09, fully corrected across all voxels and the 4 measures, not shown) in a similar area of WM (90% overlap) to that showing group differences (depicted in Fig. 2A). Animals with higher FA (and lower RD) tended to show steeper slopes (i.e. they reached criterion performance with fewer exposure days), suggesting that WM microstructure is related to TDT performance. This correlation could either reflect experience-dependent changes that occurred with task performance, or pre-existing structural differences that give rise to performance variation, or a combination of both.

### Grey Matter (GM) MD Is Lower In The TDT Group

To assess effects of experience on grey matter (GM) microstructure we tested for MD differences, as this measure reflects water restriction regardless of the structure orientation and can potentially indicate changes in tissue density and/or water content. This analysis revealed clusters with significantly lower MD (p < 0.05, corrected) in the TDT group (n=28) compared to the PC group (n=20) (Fig. 3). The significant clusters included both frontal and sensory cortex, hippocampus, and subcortical structures such as striatum and thalamic nuclei.

**Fig. 3.**
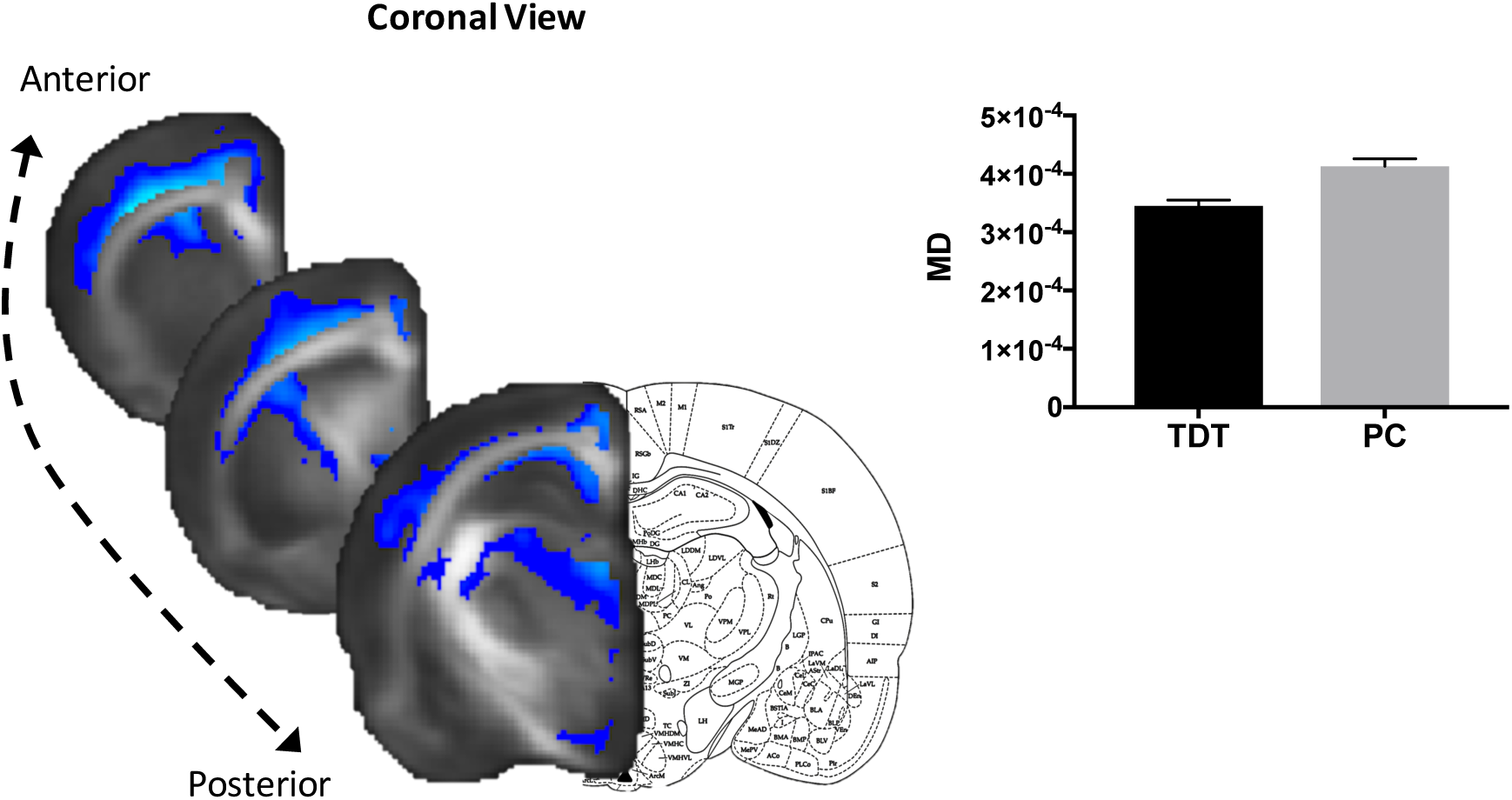
Grey Matter analysis. MD (in blue) was significantly lower in the TDT group (p < 0.05, fully corrected). Significant clusters are superimposed on the mean FA template. Bar graph of MD estimated marginal means (adjusted for the number of exposure days used as a covariate in the model) of the significant cluster is shown to illustrate the direction of differences and not for inference. Error bars represent standard error.

### Learning Versus Experience

Our main analysis above compared the TDT group to a passive control group in order to identify general experience-dependent effects. To test for specific effects of associative learning versus experience in WM microstructure, we compared a subgroup of TDT rats (TDTsg, n = 12) to an active control group (AC) (n = 12). The AC group was matched for number of training days to a subgroup of TDT rats and were exposed to the same texture discrimination apparatus but were provided with rewards that were not contingent on their response. This allowed the AC group to experience the same textures and similar levels of rewards but without the requirement to differentiate between rough and smooth textures in order to gain the rewards. Rats in TDTsg and AC groups spent the exact same time in the task (for both groups, mean = 8.83 days, S.D. = 3.97).

We used the same statistical approach as above and performed a NPC Fisher’s joint inference analysis as implemented by PALM, over the 4 DTI measures across all voxels in the WM skeleton, using the Fisher’s combining function (Winkler et al., 2016). We tested for differences between the TDTsg (n=12), AC (n = 12) and PC (n=20) groups.

The NPC joint inference test did not reveal significant differences between these groups when considering all measures together. However, the partial tests revealed several trends for FA. A trend was seen for higher FA in the AC group compared to the PC group in a cluster underlying barrel cortex (p = 0.1, fully corrected, Fig. 4A, C) and in a very similar cluster for the TDTsg group compared to the PC group (p = 0.09, fully corrected, Fig. 4B, C). There were no significant differences nor trends between the TDTsg and AC groups (p = 0.71, fully corrected). These results suggest that learning to distinguish between smooth versus rough textures is not necessary for the detected structural differences and that exposure to the task apparatus, whisker stimulation, and rewards, is sufficient to elicit similar structural changes in both active groups compared to a caged group. This indicates that the requirement to differentiate between rough and smooth textures is not necessary to elicit structural WM changes.

**Fig. 4.**
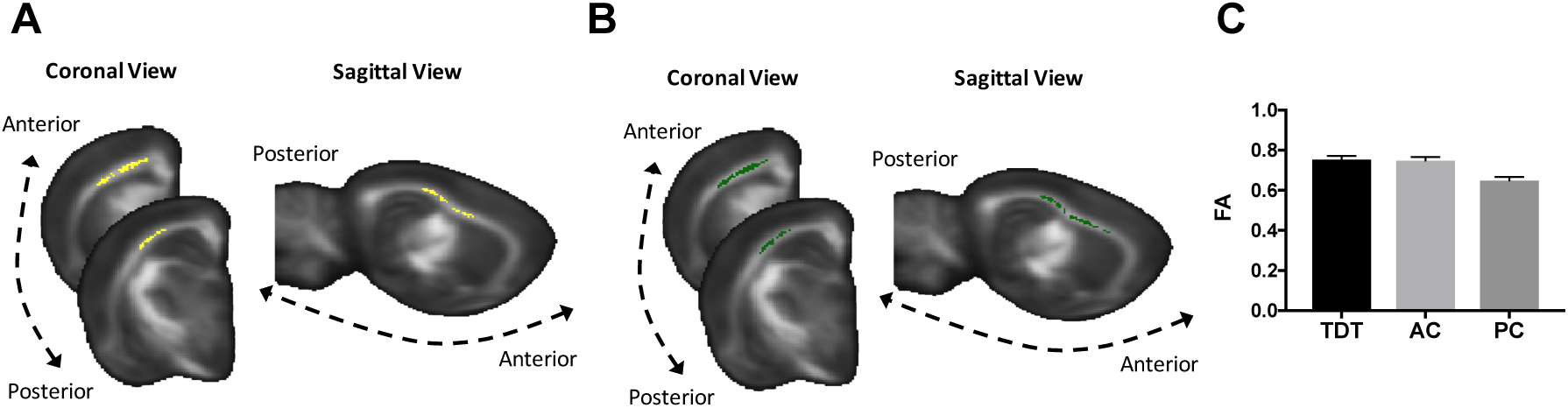
Effects of learning and experience in WM. **A** NPC partial tests revealed a trend for higher FA (yellow cluster) in the AC group compared to the PC group (p = 0.1, fully corrected). **B** NPC partial tests revealed a trend for higher FA (green cluster) in the TDTsg group compared to the PC (p = 0.09, fully corrected). No differences or trends were found between the TDTsg group and the AC. **C** Bar graph of FA estimated marginal means (adjusted for the number of exposure days used as a covariate in the model) of the overlapping cluster areas illustrated in A and B. This is shown to illustrate the direction of differences and not for inference. Error bars represent standard error.

## CANDIDATE GENE ANALYSIS RESULTS

### Synaptic C-Fos mRNA Expression In The Barrel Cortex Is Higher In The TDT Group

We assessed synaptic C-FOS expression as an indirect marker of cell activity in the barrel cortex to confirm activation of this area in response to the task. As expected, c-FOS mRNA expression was found to be higher in the TDT group compared to the PC (F_(1,23)_=11.346; p < 0.01) (Fig. 5A).

**Fig. 5.**
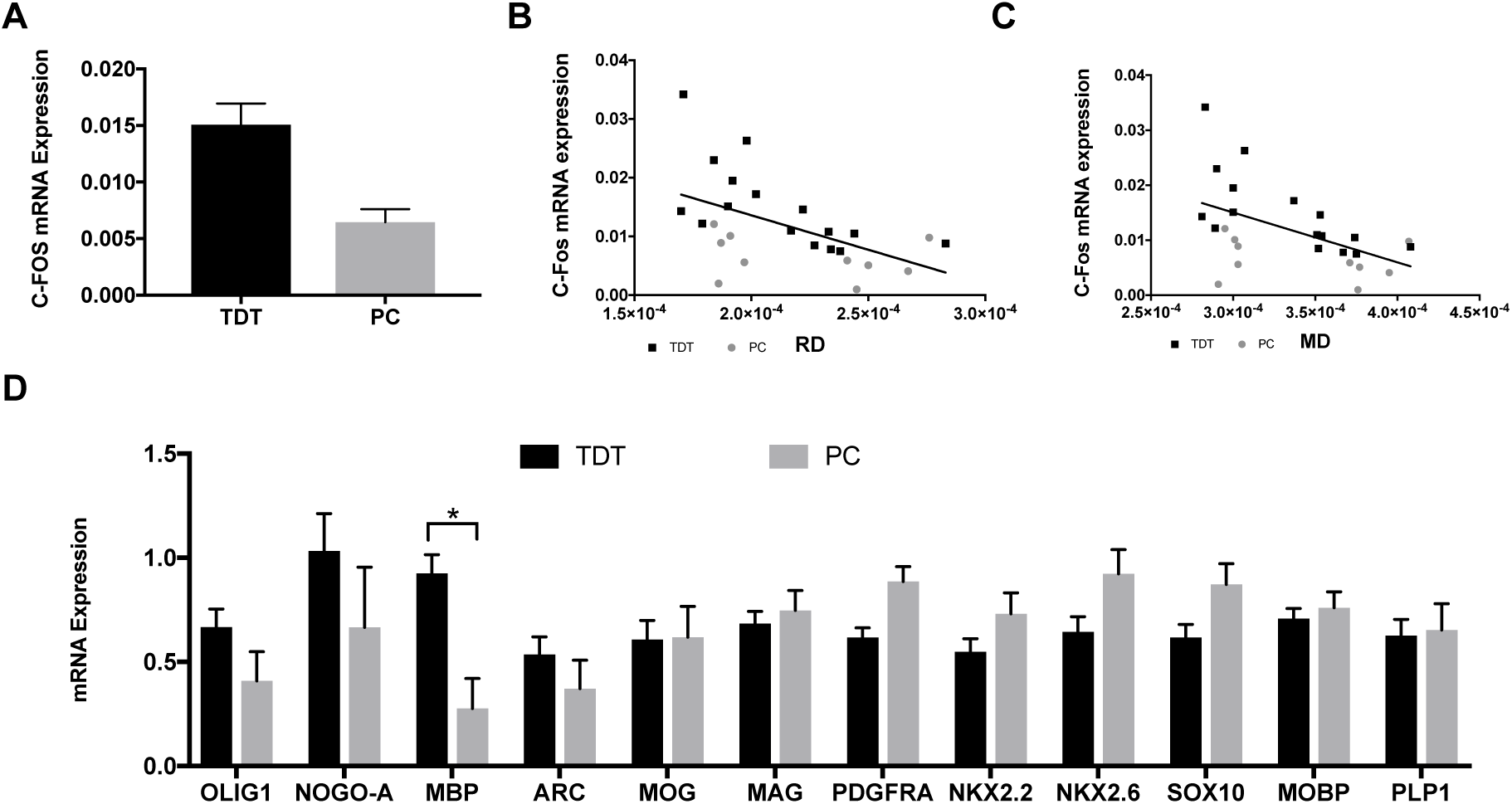
Candidate genes mRNA analysis. **A** C-FOS mRNA expression levels of the barrel cortex. **B** Plot of the significant correlation between c-FOS mRNA expression and mean RD of the significant cluster identified in Fig. 2A. **C** Plot of the significant correlation between c-FOS mRNA expression and mean MD of the significant cluster identified in Fig. 2A. **D** MRNA expression levels of myelin-related candidate genes in WM tissue underlying S1; MBP mRNA expression is higher in the TDT group compared to the control group (p < 0.05, corrected for multiple comparisons). TDT is represented in black and PC is represented in grey across all graphs.

### Synaptic C-Fos mRNA Expression In The Barrel Cortex Correlates With DTI Measures Of WM Microstructure

We further tested for correlations between c-FOS mRNA and the mean FA, MD and RD values of the significant cluster identified in Fig. 2A. We used Bonferroni correction accepting a p-value smaller than 0.0083 as significant.

There were significant negative correlations (Fig. 5B, C) across both groups (TDT and PC) between c-FOS mRNA expression and the mean RD (Pearson r = − 0.52, p = 0.006, 2-tail), and mean MD (Pearson r = − 0.51, p = 0.0078, 2-tail) but not with the mean FA after Bonferroni correction (although a trend for a positive correlation was observed, Pearson r = 0.44, p = 0.024, 2-tail).

Significant negative correlations within the TDT group only were also significant between c-FOS mRNA expression and mean RD (Pearson r = − 0.66, p = 0.005, 2-tail), and mean MD (Pearson r = − 0.69, p = 0.003, 2-tail) (represented in black in Fig. 5B, C).

### Myelin basic protein (MBP) mRNA Expression Is Higher In The WM Co-Localised With The Barrel Cortex Of The TDT Group

To assess effects of experience on mRNA expression in the WM, the WM underlying the barrel cortex was dissected from a subset of right-brain hemispheres (n=24; 15 from the TDT group and 9 from the control group) and qPCR was performed on this tissue. Expression of 12 candidate genes known to be centrally involved in myelination was analysed with non-parametric permutation testing as implemented by PALM (Winkler et al., 2016). Of the 12 candidate genes analysed only MBP survived multiple comparisons correction. MBP mRNA expression was found to be significantly higher in the TDT group (p < 0.05, corrected for multiple comparisons, Fig. 5D).

We tested for correlations between mRNA and the mean FA, MD and RD values of the cluster found to be significant in Fig. 2A with Pearson correlation coefficient. No significant correlations were found between the mRNA expression of the candidate list and the DTI measures.

## GENOME-WIDE RNA SEQUENCING RESULTS

To gain further insight into the molecular mechanisms underlying the observed structural WM and candidate gene expression differences, we performed an unbiased genome-wide analysis of mRNA expression in WM underlying the barrel cortex, by means of RNA-sequencing in a subgroup of samples (n=19; including 5 PC, 8 TDT animals and 6 AC).

The TDT versus PC comparison led to the identification of 134 differentially expressed (DE) genes, of which 65 were up and 69 downregulated (likelihood ratio test, p ≤ 5E-6, FC cut-off |1.25|, RPKM cut-off 6) (Fig. 6A, Supplementary Table 1). From this list of 134 DE genes, 124 genes were also differentially expressed between PC and AC, in the same direction (up or down) and with similar significance as in the PC versus TDT comparison. The TDT versus AC comparison lead to only 6 DE genes (Notch3, Tns1, Zbtb16, Nxn, Yap1, Cfap43), all of them downregulated (Supplementary Table 1).

**Fig. 6.**
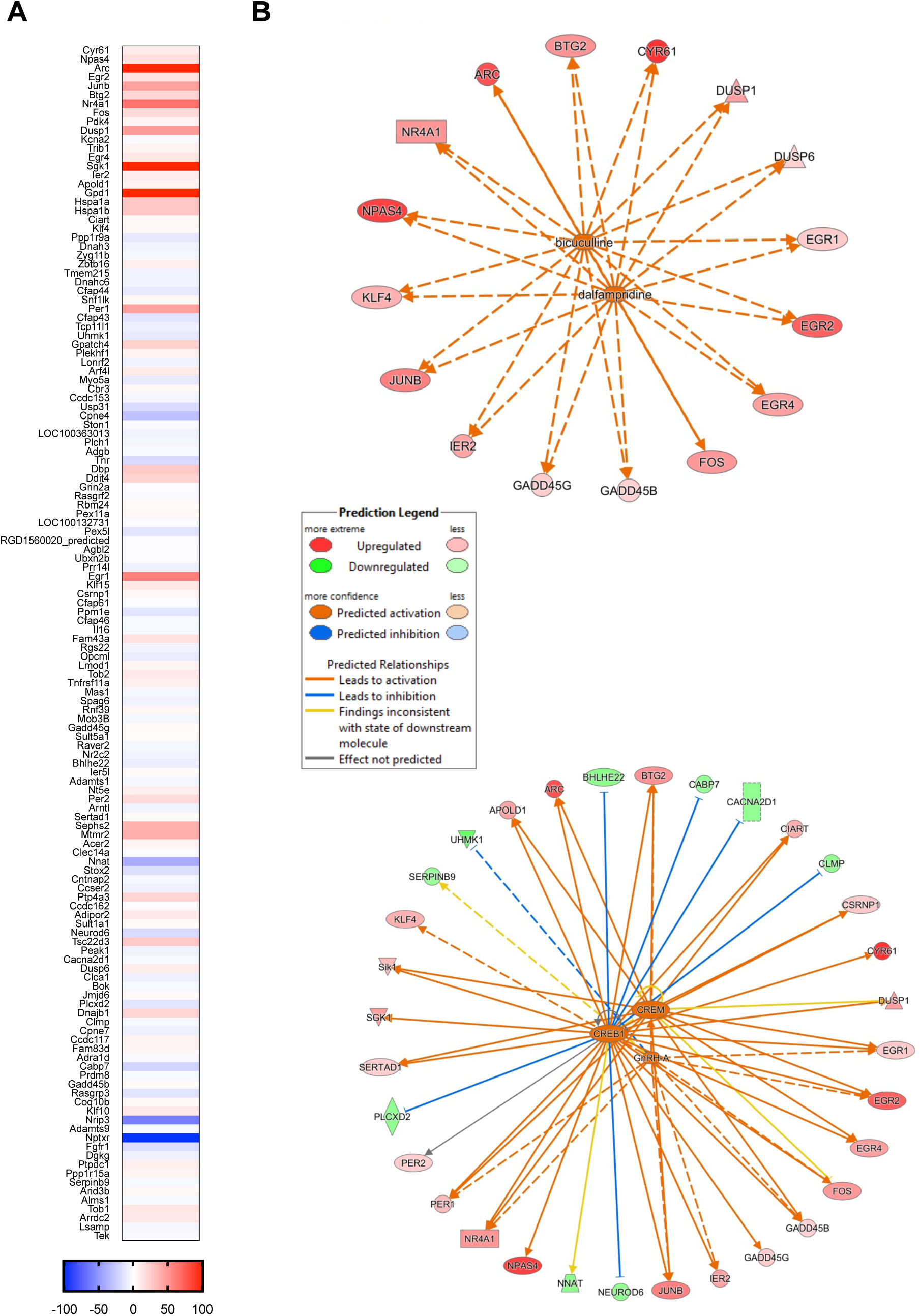
Genome-wide RNA sequencing results. **A** Heat map of 134 significantly differentially-expressed genes in TDT versus PC animals found with RNA sequencing. 124 genes of this list were also significantly differentially expressed between PC and AC. Warm colours indicate significantly upregulated (65 genes) and cold colours indicate significantly downregulated (69 genes). **B** Upstream regulators networks obtained with Ingenuity Pathway Analysis (IPA). Dalfampridine and bicuculline upstream regulator network (top). CREB1, CREM, GnRH-A upstream regulator network (bottom). Relationships between putative upstream regulators and downstream differentially-expressed molecules from the RNA sequencing dataset are shown by lines, with solid lines indicating direct relationships, and dashed lines indirect relationships. Colour coding of the lines indicates degree of concordance between predicted and actual direction of regulation. Orange lines indicate predicted activation from the upstream molecules matching observed upregulation of corresponding downstream molecules (red nodes); blue lines indicate predicted inhibition from the upstream molecule matching observed downregulation of the corresponding downstream molecule (green nodes); yellow lines indicate an inconsistent relation of the upstream regulator with the state of the downstream molecule; grey lines represent relationships not predicted by the model. Overall, there is a high concordance (few yellow lines and no grey lines) between the predicted and actual direction of regulation of the target molecules by these 5 upstream regulators.

### Gene ontology and Ingenuity Pathway analysis

In order to interpret the biological significance of the differentially expressed genes, gene ontology (GO) analysis and Ingenuity Pathways Analysis (IPA) were performed. The list of 6 DE genes (TDT versus AC comparison), led to no significant findings at the designated threshold (see Methods). In contrast, the list of 134 DE genes (TDT versus PC comparison) yielded several significant findings that are reported in detail below.

### GO analysis: MAPK signalling pathway and transcription regulator activity were enriched

GO analysis identified two significantly enriched terms (corrected p-value ≤ 0.01 (Benjamini correction) and at least 5 genes represented in the GO term were considered to be overrepresented): ‘MAPK signalling pathway’ (with 11 DE genes) and transcription regulator activity (20 DE genes) (Supplementary Table 2). MAPK signalling pathway has been implicated in cell proliferation, differentiation and development and in myelin sheath regulation (Zhang and Liu, 2002; Ishii et al., 2012).

### IPA analysis: Upstream Regulators and Networks

To identify molecules upstream of the genes that potentially explain the observed 134 DE genes, an IPA ‘Upstream Regulators’ analysis was performed (see Methods) (Supplementary Table 3). In simple terms, this analysis uses the IPA database to identify upstream regulators that match the direction of regulation of the downstream differentially-expressed molecules from the RNA sequencing dataset (Fig. 6A).

This revealed five predicted upstream regulators showing a high degree of concordance between predicted (by the IPA database) and actual direction of regulation (CREB1, CREM, GnRH-A, dalfampridine and bicuculline) (relationships illustrated in Fig. 6B), all of them with a predicted activated state, an activation z-score > 2.5 and a p-value < 5E-15, with at least 12 target molecules of the DE list dataset.

CREB1 and CREM had overlapping target molecules (29 target molecules for CREB1, of which 16 were also CREM targets), and had a high degree of overlap with GnRH-A targets (12 targets, of which 9 overlapping with CREB1/CREM targets) (Fig. 6B bottom). CREB and CREM are transcription factors activated by phosphorylation in response to cAMP and other signals (for review see (Mayr and Montminy, 2001)). Both CREB and CREM are involved in regulating the transcription of several genes (c-fos, BDNF, etc) and have been implicated in neuronal plasticity and memory (Lonze and Ginty, 2002; Benito and Barco, 2010). More recently, CREB has been connected with the transcriptional control of MBP (Meffre et al., 2015), also found to be differentially expressed in our study.

Additionally, two drugs, dalfampridine and bicuculline, shared the same 16 target molecules (Fig. 6B top). Dalfampridine is a broad-spectrum voltage-gated potassium channel blocker that broadens the action potential. It shows beneficial effects in multiple sclerosis patients, probably through restoration of axonal conduction (Dunn and Blight, 2011) and has been shown to promote remyelination after acute nerve injury (Tseng et al., 2016). Bicuculline blocks GABA_A_-mediated inhibition, thereby increasing neuronal activity. This shows that genes involved in neuronal activity modulation are differentially expressed in our sample, possibly indicating increases or functional changes in axonal activity.

Next we sought to identify functionally related networks of genes and important regulatory hubs (Fig. 7). There were two networks with a score >30 containing at least 17 molecules of the DE dataset. In Network 1, two main ‘hub’ molecules (i.e. where most relationships converge to), Akt and Creb (Fig. 7A), were identified, while in Network 2 the main ‘hub’ molecules were Erk1/2 (Fig. 7B). Erk1/Erk2, together with Akt/mTOR, have been proposed as two main signalling pathways for the control of proliferation and differentiation in oligodendrocyte precursor cells (OPCs) and in myelin sheath regulation in adult oligodendrocytes (Cui and Almazan, 2007; Bibollet-Bahena and Almazan, 2009; Dai et al., 2014; Snaidero et al., 2014).

**Fig. 7.**
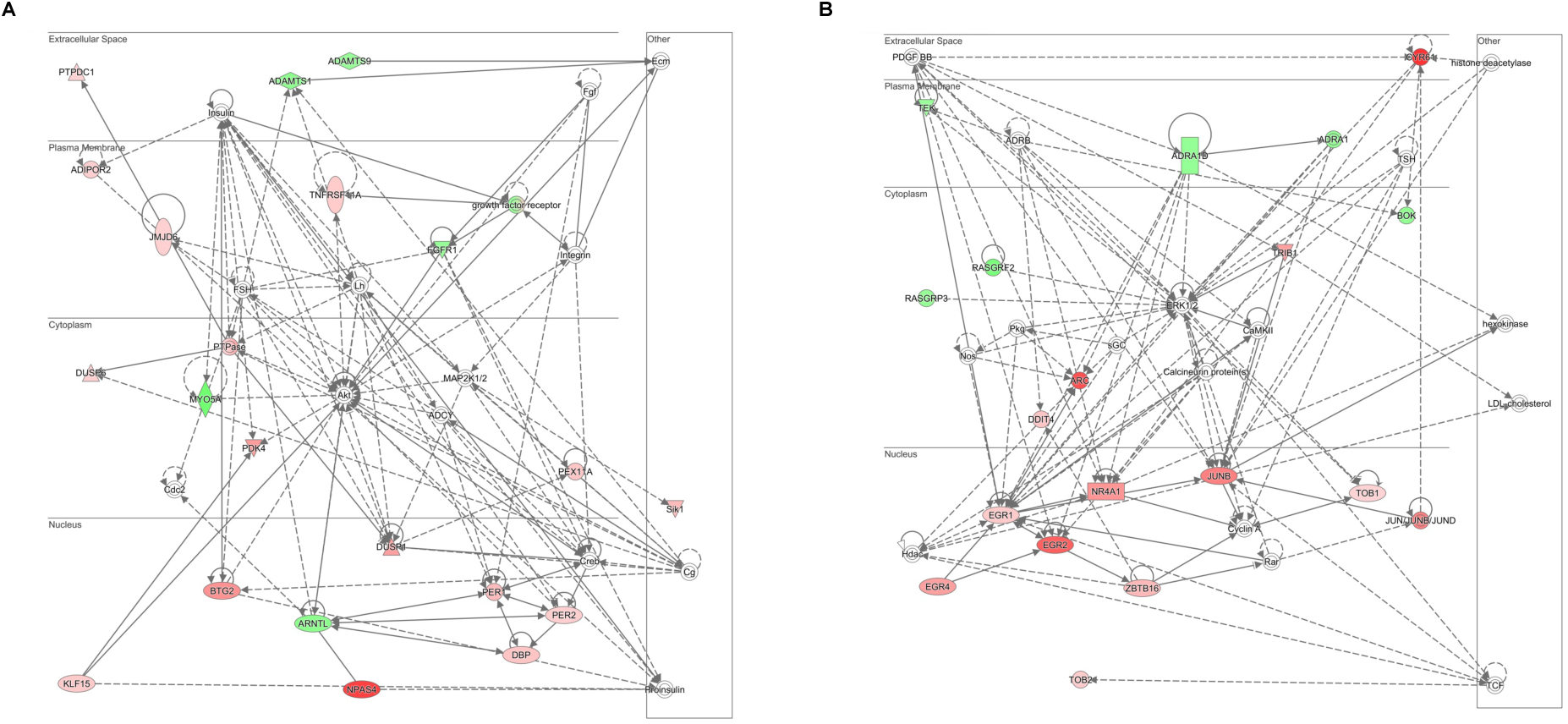
Two main networks were identified with Ingenuity Pathway Analysis (IPA) ***A*** *Network 1 with Akt and Creb as hub molecules (where most relationships converge to)* ***B*** *ERK1/2 as hub molecules in Network 2*. The upregulated DE molecules from the RNA sequencing analysis are represented in red while the downregulated are labelled in green.

## DISCUSSION

Somatosensory experience results in structural changes including increases in dendritic spines (Knott et al., 2002) and cortical myelinogenesis (Hughes et al., 2018). Here we have demonstrated that somatosensory experience also results in structural white matter plasticity and identify some molecular correlates that provide candidate mechanisms underlying these findings, such as myelin formation and/or remodelling.

Our results suggest that learning the associative task is not necessary for the detected plastic changes and that mere exposure to somatosensory stimulation is sufficient since structural or genome-wide mRNA expression differences between the TDT group and an active control group were not identified.

The WM structural diffusion metrics were found to correlate with barrel cortex synaptic c-fos expression, suggesting that molecular correlates of cortical activity relate to macroscale measures of WM structural plasticity. Synaptoneurosomes are enriched in synaptic terminals (pre and post) and might also include other cellular (for instance astrocytic) components. Although c-fos is often regarded as an immediate early gene with exclusive expression in neurons, its expression has also been shown in astrocytes under certain conditions, so we cannot completely exclude non-neuronal contribution to the synaptic c-fos expression (Herrera and Robertson, 1996). There is increasing evidence from *in vitro* (Demerens et al., 1996) and *in vivo* (Mensch et al., 2015; Etxeberria et al., 2016; Dutta et al., 2018; Piscopo et al., 2018) studies that neuronal activity modulates myelination, even in adulthood (Gibson et al., 2014). In line with this, our mRNA expression analysis indicated that myelination-related genes (e.g. MBP, Akt and Erk1/Erk2 gene networks) (Ishii et al., 2012; Jeffries et al., 2016) are differentially expressed in response to somatosensory experience.

The DTI analysis revealed higher FA and lower RD in the TDT group indicating that water diffusion is more hindered across WM tracts after somatosensory experience. Additionally, lower MD was found diffusion in both WM and GM, indicating higher overall restriction of water which could potentially be related to greater tissue density in these areas. While definitive biological interpretation of DTI changes is challenging (Sampaio-Baptista and Johansen-Berg, 2017), this pattern of DTI differences in WM is consistent with cellular mechanisms such as higher myelin thickness or internode length (Etxeberria et al., 2016). Accordingly, we found higher MBP mRNA expression suggesting that myelination has been triggered by somatosensory experience. Increases in myelin are consistent with the higher FA and decreased RD found in the current study. These findings are congruent with previous neuroimaging studies that have found higher FA, along with higher MBP immunostaining intensity in response to complex motor and cognitive learning (Blumenfeld-Katzir et al., 2011; Sampaio-Baptista et al., 2013). However, additional mechanisms may also contribute to these findings. For example, changes in axon diameter (Sinclair et al., 2017), nodes of Ranvier length (Arancibia-Carcamo et al., 2017) or axon packing density can potentially also be reflected in these DTI measures. Given the large number of astrocytes present in WM, alterations in this cell population can also potentially modulate DTI measurements (Sampaio-Baptista and Johansen-Berg, 2017). For instance, MD decreases and astrocyte changes have been described in GM in learning paradigms (Blumenfeld-Katzir et al., 2011; Johansen-Berg et al., 2012; Sagi et al., 2012), but there is currently very little understanding of structural contributions of astrocytes to the diffusion signal in the context of long-term experience in WM. Evaluation of the volume, shape and size of astrocytes using immunohistochemistry or other techniques after experience or learning paradigms along with DTI measures would help to clarify in which direction to formulate predictions.

The genome-wide mRNA analysis identified 134 differentially expressed genes that are associated with functions related to neuronal plasticity (CREB1, CREM), memory (CREB1, CREM), neuronal activity modulation (16 target molecules of voltage-gated potassium channels and GABA_A_-mediated inhibition blockers), myelin-sheath thickness regulation (Erk1/Erk2), and proliferation control, differentiation and protein synthesis in OPCs (CREB1, CREM, Akt and Erk1/Erk2). Together these indicate that WM has undergone functional and structural plasticity in response to somatosensory experience. In particular, Erk1/Erk2, together with Akt/mTOR, are the two main signalling pathways for the control of proliferation, differentiation and protein synthesis in oligodendrocyte precursor cells (OPCs) (Cui and Almazan, 2007; Bibollet-Bahena and Almazan, 2009; Dai et al., 2014). Akt and mTOR signalling pathway plays a central role in promoting myelination (reviewed in (Norrmen and Suter, 2013)) and forced activation of the Akt pathway in adult oligodendrocytes results in growth of myelin sheaths (Snaidero et al., 2014). Erk1/Erk2, from the mitogen-activated protein kinase (MAPK) pathway (also significantly enriched in our analysis), is also an important regulator of myelin-sheath thickness in the CNS (Ishii et al., 2012; Jeffries et al., 2016). Conditional upregulation of Erk1/Erk2 results in global increases in myelin thickness by preexisting oligodendrocytes of adult mice, faster nerve conduction velocity and behavioural changes (Jeffries et al., 2016). Furthermore, Erk2 has been described to have an important role in oligodendrocytes in the translational control of MBP (Michel et al., 2015), which is also in line with our findings. Our results indicate that somatosensory experience may trigger both de novo myelin formation as well as increases in thickness of myelin sheaths.

In conclusion, somatosensory experience resulted in macroscale structural changes detected with DTI that are consistent with higher myelination, and further supported by molecular evidence showing higher MBP mRNA expression and differentially expressed genes involved in regulation of proliferation and differentiation in OPCs and in myelin sheath formation. Additionally, WM structure correlated with cortical activity as measured by c-fos mRNA, consistent with the idea that cortical experience-dependent mechanisms could trigger WM plasticity. Taken together our results demonstrate that myelination occurs in response to somatosensory experience and that this experience-dependent myelin plasticity is reflected in DTI metrics in WM.

This work paves the way for future studies to examine the specific effects of the identified genes on MRI measures by combining genetic (McKenzie et al., 2014; Jeffries et al., 2016) or pharmacological manipulation in rodents with imaging read-outs. This would allow to precisely identify the molecular and cellular mechanisms which underlie changes in MRI measures of plasticity and could offer important clues to the biological changes underlying imaging signals recorded in humans.

## MATERIAL AND METHODS

### Animals

All behavioural experiments were conducted at Radboud University Nijmegen (The Netherlands). The experiments were approved by the Animal Ethics Committee of the Radboud University Nijmegen (The Netherlands), according to Dutch legislation and all procedures were performed according to the project and personal licenses held by the experimenters.

60 animals (3 months old, male Long Evans rats (250 - 450 g) (Harlan, Bicester, UK)) were housed in standard laboratory conditions under a 12-h light/12-h dark cycle at 20 °C temperature and 40 – 70 % humidity. The animals were housed individually for more precise control of their general welfare and because group housing may interfere with the task experience. All animals were given appropriate time to acclimate after delivery (1 week minimum) and had ad libitum access to food and water. After this period, they were handled daily for one week before the start of the task. Before the task, animals were exposed to the testing arena for 10 minutes each day for 1-2 days under dim visible light.

The animals were given no access to water for a period of 24h before the first session. From here on, they received water during the task (0.1 ml of water per correct trial) and water was also made available ad libitum for 30 min after the session. The delay between the end of the task and the time period when water was freely available varied between 30 minutes to 2 hours in order to prevent the animals from learning that water would be available after the testing period.

### Texture Detection Task

The texture detection task (TDT) is based on a previously described task (von Heimendahl et al., 2007). Rats were trained to distinguish between a smooth and a textured surface using operant conditioning as described below. Training was performed in the dark to avoid the influence of visual cues on performance. Potential olfactory cues were removed from textures by washing them at least once every individual animal session, and by using different sets of identical textures that were interchanged randomly between animals and sessions.

Rats were tested individually. During testing, the animal was placed on a 30 cm elevated platform with two water dispensers on each side. Under this platform was a small bridge where the animal could place its front paws for a short period of time in order to reach the stimulus presented in front of the platform. The stimulus consisted of a series of rectangle shapes with patterns that could be varied depending on the animal’s performance in the task.

During the shaping period the animal was placed on the apparatus, and every appropriate response was rewarded. First, water was randomly delivered in order for the animal to learn where the water was placed. After the animal had learned this, it was rewarded for leaning on the edge of the platform and reaching the stimulus. Finally, water was delivered when the animal touched the texture with its whiskers.

During the training phase of the TDT, the stimulus was either a smooth texture (reference texture) or a positive copy of sandpaper on a resin material. Each animal was trained with a fixed association (e.g., turn left on rough, right on smooth). Only if it approached the correct drinking spout, the animal was given a water reward (0.1 ml per reward); for an incorrect choice, it received no water. The next trial started with a delay of 5 s. Between trials, the texture’s stand was turned about its vertical axis by a computer-controlled stepping motor, which allowed for quick, randomized, and automated switching between textures. Each session lasted for about 30 min, during which the animal performed between 60-100 trials.

### Experimental Design

Animals were randomly assigned to the TDT and control groups, balancing for weight to obtain equal weight averages between the groups.

Texture Detection Task (TDT) group: For the TDT, animals (total n=28) were trained to distinguish between the reference texture (smooth) and a P100 texture (162 um average particle diameter). Rats were trained 5 days a week. Individual rats were sacrificed the day after they reached criterion (2 sessions at > 80% accuracy) on the P100 texture, in order to have comparable performance levels between animals. This resulted in a variation in the number of task exposure days which was then controlled for in subsequent analyses as described below.

A subgroup of rats (n = 8) were further trained to detect increasingly more fine-grained textures after reaching performance criterion in the P100 texture in a stepwise manner: P150 (100 um average particle diameter), P220 (68 um), P280 (52.2 um), P360 (40.5 um), P400 (35 um), P500 (30.2 um) and P600 (25.8 um). When the animals performed above criterion (> 80% accuracy) for a given texture, the rough texture was changed to a finer one on the following training day. Rats were sacrificed the day after they reached > 80% accuracy on the P600 texture. After the rats had associated the correct reward side with the first texture, increasing the difficulty of the texture discrimination did not alter their accuracy (Supplementary Fig. 1). Negative control experiments were performed in this subgroup of rats to demonstrate that the animals distinguished smooth vs rough textures and not other sensory attributes of the task (e.g. noises or odours). To do that, animals were presented with the same texture (P400 versus P400) and their performance was assessed. When rats were presented with the same texture their performance accuracy dropped to chance levels (Supplementary Fig. 1).

#### Active Control (AC) group

12 rats were matched to an individual in the TDT group. Rats were water restricted and exposed to the TDT task for the same period of time as the matched animal. However, these animals were rewarded randomly, and not in relation to texture-response contingencies. They received a similar number of rewards as the matched animal throughout the entire training period and were sacrificed after the same number of training sessions as the matched animal in the TDT group.

#### Passive Control (PC) group

Caged controls (n = 20) were handled and weighed daily; their body weight served as a reference body weight with respect to the other group.

### Brain Preparation

TDT rats were sacrificed by rapid decapitation without anaesthesia on the day after they reached criterion. The AC group were sacrificed after the same number of training sessions as the matched animal in the TDT group. On the day of sacrifice, animals were trained on their respective task for 15 minutes, then placed back in their home cage for 15 minutes, after which they were sacrificed and the brains were removed. The PC group was handled for 15 min prior to the sacrifice. The right hemisphere was frozen on dry ice and kept at −80 °C for molecular analysis and the left hemisphere was immersed in 4% PFA for DTI acquisition.

For DTI acquisition, all left brain hemispheres (n=60) were placed into falcon tubes (50ml) in pairs (one from each group), one hemisphere above the other, and embedded in 2 % agarose gel (Sigma-Aldrich) (Sampaio-Baptista et al., 2013). The hemispheres were aligned to each other along the posterior – anterior axis.

### MRI Acquisition

All 60 ex-vivo left brain hemispheres were scanned in pairs overnight with a 4.7T MRI scanner (Agilent Technology Inc., USA) at Radiobiology Research Institute, Churchill Hospital, Oxford. DTI scanning parameters were as follows: Spin-Echo Multi-Slice Diffusion Weighted (SEMSDW) sequence, b = 2000 s/mm^2^, 30 diffusion directions, 4 averages plus 8 images with no diffusion weighting, 40 slices, slice thickness 0.5 mm, field of view 25×50 mm, matrix size 96 × 192 (resolution 0.26 × 0.26 × 0.5 mm).

### Tissue Dissection

A subset of right brain hemispheres was dissected for qPCR and RNA-SEQ. The experimenter was blind to the group for all the following procedures. All procedures were performed under RNase-free conditions. The brain hemispheres were sliced into 300 µm coronal sections using a cryotome (Leica GmbH, Germany) at −15 °C and mounted on glass slides. Cytochrome oxidase-stained reference sections were used as a template to locate the barrel cortex, following stereotactic coordinates (Paxinos and Watson, 1998). Punches of the barrel cortex (n=26; 16 from the TDT group, 10 from the PC group) and in WM (n=30; 15 from the TDT group, 6 from the AC, 9 from the PC group) directly underneath it of the right hemisphere were taken using a 1.20-mm micropunch (Harris Inc., UK) and stored at −80 °C before RNA isolation took place.

### RNA Isolation

The experimenter was blind to the group in all the following procedures and the groups were randomly distributed to ensure equal distribution of groups to avoid any technical bias. Samples were homogenized with a TissueLyser (Retsch GmbH, Germany) in TRIzol® Reagent (Invitrogen Co., USA). RNA was isolated with TRIzol® Reagent (Invitrogen), according to the manufacturers’ protocol. The procedure was modified for small amounts of tissue by using 800 μl of TRIzol® Reagent and adding 1 μl of glycogen (Fermentas Inc., USA). RNA concentration and quality was determined with a NanodropTM ND-1000 spectrophotometer (Thermo Fisher Scientific Inc., USA) and 1% agarose gel electrophoresis, respectively. The samples were kept at −80 °C until further analysis.

### Synaptoneurosome Preparation

Synaptoneurosomes were prepared by the method described by (Williams et al., 2009), with some modifications. Brain tissue punches were homogenized with a Teflon-homogenizer (12-14 strokes at 1000 rpm) in 4 mL of homogenization buffer, containing 0.35 M sucrose pH 7.4, 10 mM 4-(2-hydroxyethyl)-1-piperazineethanesulfonic acid (HEPES), 1 mM ethylenediaminetetraacetic acid (EDTA), 0.25 mM dithiothreitol, 8 U/ml RNAse inhibitor and a protease inhibitor cocktail (Roche). Cell debris and nuclei were removed by centrifugation at 1000 g for 10 min at 4°C yielding pellet P1 and supernatant S1. The S1 fraction was passed sequentially through a series of filters with decreasing pore sizes of 80, 40 and 10 µm (Millipore). The final filtrate was centrifuged at 2000g for 15min at 4°C yielding pellet P2 and supernatant S2. Pellet P2 containing synaptoneurosomes was resuspended in 200 µL of homogenization buffer. Enrichment of synaptic components in the synaptoneurosomal fraction was assessed by western blot in control experiments. To assess c-fos expression in synaptoneurosomes, RNA was isolated using the Trizol method, followed by downstream QPCR.

### Quantitative PCR (qPCR)

Prior to cDNA-synthesis, 0.5 μg of each RNA sample was treated with 2 U DNase (Fermentas Inc., USA), in the presence of RiboLockTM RNase Inhibitor (20 U/μl) (Fermentas Inc., USA). For cDNA synthesis, through random priming, the RevertAidTM H Minus First Strand cDNA Synthesis kit (Fermentas Inc., USA) was used, following the manufacturer’s guidelines. Prior to analysis, each cDNA sample was diluted 1/15 with MilliQ water. qPCR reactions were performed with the Rotor-Gene 6000 Series (Corbett Life Science Pty. Ltd., Australia). For each reaction, 2.5 µL of each diluted sample of cDNA was added to a mix containing 6.25 µL 2X MaximaTM SYBR Green qPCR Master Mix (Fermentas Inc., USA), 1 µL of each primer (5 μM) and 1.75 μL MilliQ water. Primers were designed using NCBI Primer-Blast (www.ncbi.nlm.nih.gov/tools/primer-blast/) and synthesized at Sigma-Aldrich (UK). Cycling conditions were 10 min 95°C followed by 40 cycles of 15 sec at 95°C, 30 sec at 60°C and 30 sec at 72°C. After cycling, a melting protocol was performed, from 72 °C to 95 °C, measuring fluorescence every 1 °C, to control for product specificity.

For the candidate gene analysis of the WM, the following genes were selected for their role in myelin and WM plasticity (PLP1, OLIG1, NOGO-A, MBP, MOG, MAG, PDGFRA, NKX2.2, NKX2.6, SOX10, MOBP, ARC). The c-FOS gene was selected to confirm Barrel Cortex neuronal activation in response to the TDT task. Relative expression of the selected genes of interest was calculated after obtaining the corresponding Ct values and correcting for unequal sample input using geNorm (Vandesompele et al., 2002), which identifies the two most stably expressed housekeeping genes from a set of three tested candidate genes (ACTB, YWHAZ and CYCA) previously reported to be stably expressed in the brain (Bonefeld et al., 2008) to calculate a normalization factor for each sample. This normalization factor was then used to obtain the relative differences between the samples for each primer pair.

### RNA Sequencing (RNA-Seq)

A subgroup of samples (n=13; 5 PC, 8 TDT animals and 6 AC) was used for RNA-Seq, four pools of total RNA (with equal input from each individual sample) were made after Trizol extraction, and were further purified and DNase-treated using Qiagen colums (RNeasy Plus Microkit, Quiagen). The yield of the purified RNA ranged between 1.5 and 2 ug total RNA per pool. The five pools were as follows: (1) passive control (PC) (n=5 individual samples in pool), (2) TDT Long-exposure (LE) (n=2 individual samples in pool), (3) TDT Mid-exposure (ME) (n=3 individual samples in pool), (4) TDT Short-exposure (SE) (n=3 individual samples in pool) and (5) active control (AC) (n=6 individual samples in pool). The division of the TDT group into 3 pools of samples was made on basis of the number of days that the animals needed to reach criterion in the behavioural experiment: LE: 9-10 days, ME: 6-7 days, SE: 5 days.

The purified RNA pools were sent for further quality control and RNA-Seq analysis to the Genomic Services Lab of the HudsonAlpha Institute for Biotechnology (AL, USA); http://gsl.hudsonalpha.org/. RNA-Seq with ribosomal RNA (rRNA) reduction was used using standard protocols (depth >45M pair-end reads per sample), and the resulted raw data were received. Data analyses (alignment and statistical analysis) were performed with GeneSifter (Geospiza, PerkinElmer Inc). We compared the PC group with all three TDT groups pools (LS, LM, LF). For each comparison a p value (likelihood ratio test) and fold change (FC) was obtained and the following cut-offs were applied: p value ≤5E-6 in all 3 (PC vs LS, LM, LF) comparisons, FC>|1.25| in at least 2 out of 3 comparisons (with a minimum FC>|1.2|), and expression levels (reads per kilobase million, RPKM) > 6 in at least 1 out of the 4 groups compared. Additionally, to test for specific effects of associative learning versus experience the AC group was also compared to the TDT groups pools and the same cut-offs described above were applied.

### MRI Statisitical Analysis

The data were pre-processed according to standard procedures in FSL (Smith et al., 2004). Tract Based Spatial Statistics (TBSS) (Smith et al., 2006) was applied to the pre-processed data. Images were then analysed as described elsewhere (Sampaio-Baptista et al., 2013). Briefly, all FA maps were aligned with linear and non-linear transformations to the study specific template and averaged to generate the mean FA image, from which the WM skeleton was extracted. The skeleton was thresholded at an FA value of 0.36 to contain only the major tracts (Sampaio-Baptista et al., 2013). Finally, the FA values of the tract centres were projected onto the skeleton for each rat brain and fed into statistical analysis. MD, RD and AD skeleton maps were created with the same method, using the FA registrations and skeleton projections as implemented in TBSS for non-FA images (Smith et al., 2006).

We used Permutation Analysis of Linear Models (PALM) (Winkler et al., 2016) for multi-measures analysis. PALM is a tool that allows inference over multiple modalities, including non-imaging data, using non-parametric permutation methods, similarly to the *randomise* tool in FSL (Winkler et al., 2014), although offering a number of features not available in other analysis software, such as the ability for joint inference over multiple modalities, or multiple contrasts, or both together, while correcting FWER or FDR across modalities and contrasts (Winkler et al., 2016).

We used PALM to assess the joint and individual contribution of the 4 DTI measures while simultaneously correcting across the tests. Non-Parametric Combination (NPC), as implemented in PALM, was used for joint inference over the 4 DTI measures (FA, MD, RD and AD). NPC works by combining test statistics or p-values of separate (even if not independent) analyses into a single, joint statistic, the significance of which is assessed through synchronized permutations for each of the separate tests. The synchronized permutations for the separate tests accommodate, implicitly, any eventual lack of independence among them. Such a joint analysis can be interpreted as a more powerful, permutation-based version of the classical multivariate analysis of covariance (MANCOVA); differently than MANCOVA, however, NPC allows investigation of the direction of joint effects.

Here we used NPC with Fisher’s combining function, testing for effects with concordant directions across the 4 DTI measures. A cluster-forming threshold of t > 1.7 and 5000 permutations were used to determine p-values FWER-corrected for multiple comparisons (across all voxels and the 4 DTI measures). The chosen cluster-forming t threshold was based on the degrees of freedom of the sample. Clusters with a corrected significance of p < 0.05 were deemed significant.

We performed two statistical tests in the WM analysis. First we tested for differences between groups and included the total number of exposure days per animal as a covariate. Second, we tested for correlations between performance rate and the 4 DTI measures. This was calculated by fitting a logarithmic model and extracting the slope of the percentage of correct trials curve for each individual animal (curves are illustrated in Fig. 1A).

Further, we tested for GM differences using MD only as this measure can indicate changes in tissue density regardless of the structure orientation. We tested for group differences and included the total number of exposure days per animal as a covariate. We performed non-parametric permutation testing with the Randomise tool as implemented in FSL (Smith et al., 2004), with a cluster-forming threshold of t > 1.7 and 5000 permutations were used to determine corrected p-values. Clusters with a corrected significance of p < 0.05 were deemed significant.

### Statistical Analysis of mRNA Expression

Statistical analysis of WM qPCR data was also performed with PALM (Winkler et al., 2016). We tested for group differences with non-parametric permutation testing with a between groups contrast (with total number of exposure days as a covariate). A p-value of < 0.05 was deemed significant, corrected for multiple comparisons across the 12 genes of interest.

For the qPCR analysis of the c-FOS gene of the Barrel cortex, a one-way analysis of covariance was used to test for differences between groups (with total number of exposure days as a covariate) with SPSS. A p-value of < 0.05 was deemed significant.

### Gene Ontology (GO) Enrichment Analysis and Ingenuity Pathways Analysis (IPA)

Gene Ontology (GO) enrichment analysis of the differentially expressed genes was performed using the web-based gene ontology tool from the Database for Annotation, Visualization and Integrated Discovery (DAVID) 6.7 (http://david.ncifcrf.gov) (Dennis et al., 2003; Huang da et al., 2009b, a). For the enrichment analysis (Functional Annotation Chart tool), default software settings were used, and GO terms with a corrected p-value ≤ 0.01 (Benjamini correction) and at least 5 genes represented in the GO term were considered to be overrepresented (enriched).

Ingenuity Pathways Analysis (IPA) (Ingenuity Systems Inc., USA), was used to perform pathway, network and upstream regulator analyses to explore relationships between genes on the basis of curated information present in the IPA database. For pathway and interaction network analyses, a score was obtained (calculated as the –log of the associated Fisher’s exact test p value). This score indicates the likelihood that the assembly of a set of focus genes in a network could be explained by random chance alone; networks with scores of 2 or higher have at least a 99% confidence of not being generated by random chance alone. Upstream regulator analysis generated a list of putative upstream regulators of the DE genes, and indicate, for each putative upstream regulator, a predicted activation state, activation z-score, p value of overlap and list of putative target genes of the DE dataset.

## ACKNOWLEDGEMENTS

The authors would like to thank Riejanne Seigers for her valuable help with the behavioural training, and Ivica Granic for his technical support with the tissue dissections.

This work was supported by the Wellcome Trust (WT090955AIA and WT110027/Z/15/Z to H J-B). C S-B was the recipient of a FCT fellowship (SFRH/BD/43862/2008). The Wellcome Centre for Integrative Neuroimaging is supported by core funding from the Wellcome Trust (203139/Z/16/Z). AK and NS were funded by Cancer Research UK (C5255/A15935). This research was further supported by the Dutch Science Foundation (NWO) through VICI grant 453-04-002 (to P.D.W.) and VENI grant 451-09-025 (to M.R.).

## REFERENCES

Arancibia-Carcamo IL, Ford MC, Cossell L, Ishida K, Tohyama K, Attwell D (2017) Node of Ranvier length as a potential regulator of myelinated axon conduction speed. Elife 6.

Benito E, Barco A (2010) CREB’s control of intrinsic and synaptic plasticity: implications for CREB-dependent memory models. Trends Neurosci 33:230–240.

Bibollet-Bahena O, Almazan G (2009) IGF-1-stimulated protein synthesis in oligodendrocyte progenitors requires PI3K/mTOR/Akt and MEK/ERK pathways. J Neurochem 109:1440–1451.

Blumenfeld-Katzir T, Pasternak O, Dagan M, Assaf Y (2011) Diffusion MRI of structural brain plasticity induced by a learning and memory task. PLoS One 6:e20678.

Bonefeld BE, Elfving B, Wegener G (2008) Reference genes for normalization: a study of rat brain tissue. Synapse 62:302–309.

Cui QL, Almazan G (2007) IGF-I-induced oligodendrocyte progenitor proliferation requires PI3K/Akt, MEK/ERK, and Src-like tyrosine kinases. J Neurochem 100:1480–1493.

Dai J, Bercury KK, Macklin WB (2014) Interaction of mTOR and Erk1/2 signaling to regulate oligodendrocyte differentiation. Glia 62:2096–2109.

Demerens C, Stankoff B, Logak M, Anglade P, Allinquant B, Couraud F, Zalc B, Lubetzki C (1996) Induction of myelination in the central nervous system by electrical activity. Proc Natl Acad Sci U S A 93:9887–9892.

Dennis G, Jr., Sherman BT, Hosack DA, Yang J, Gao W, Lane HC, Lempicki RA (2003) DAVID: Database for Annotation, Visualization, and Integrated Discovery. Genome Biol 4:P3.

Dunn J, Blight A (2011) Dalfampridine: a brief review of its mechanism of action and efficacy as a treatment to improve walking in patients with multiple sclerosis. Curr Med Res Opin 27:1415–1423.

Dutta DJ, Woo DH, Lee PR, Pajevic S, Bukalo O, Huffman WC, Wake H, Basser PJ, SheikhBahaei S, Lazarevic V, Smith JC, Fields RD (2018) Regulation of myelin structure and conduction velocity by perinodal astrocytes. Proc Natl Acad Sci U S A 115:11832–11837.

Etxeberria A, Hokanson KC, Dao DQ, Mayoral SR, Mei F, Redmond SA, Ullian EM, Chan JR (2016) Dynamic Modulation of Myelination in Response to Visual Stimuli Alters Optic Nerve Conduction Velocity. J Neurosci 36:6937–6948.

Feldmeyer D, Brecht M, Helmchen F, Petersen CC, Poulet JF, Staiger JF, Luhmann HJ, Schwarz C (2013) Barrel cortex function. Prog Neurobiol 103:3–27.

Gibson EM, Purger D, Mount CW, Goldstein AK, Lin GL, Wood LS, Inema I, Miller SE, Bieri G, Zuchero JB, Barres BA, Woo PJ, Vogel H, Monje M (2014) Neuronal activity promotes oligodendrogenesis and adaptive myelination in the mammalian brain. Science 344:1252304.

Herrera DG, Robertson HA (1996) Activation of c-fos in the brain. Prog Neurobiol 50:83–107.

Hofstetter S, Tavor I, Tzur Moryosef S, Assaf Y (2013) Short-term learning induces white matter plasticity in the fornix. J Neurosci 33:12844–12850.

Holtmaat A, Wilbrecht L, Knott GW, Welker E, Svoboda K (2006) Experience-dependent and cell-type-specific spine growth in the neocortex. Nature 441:979–983.

Huang da W, Sherman BT, Lempicki RA (2009a) Bioinformatics enrichment tools: paths toward the comprehensive functional analysis of large gene lists. Nucleic Acids Res 37:1–13.

Huang da W, Sherman BT, Lempicki RA (2009b) Systematic and integrative analysis of large gene lists using DAVID bioinformatics resources. Nat Protoc 4:44–57.

Hughes EG, Orthmann-Murphy JL, Langseth AJ, Bergles DE (2018) Myelin remodeling through experience-dependent oligodendrogenesis in the adult somatosensory cortex. Nat Neurosci 21:696–706.

Ishibashi H (2002) Increased synaptophysin expression through whisker stimulation in rat. Cell Mol Neurobiol 22:191–195.

Ishii A, Fyffe-Maricich SL, Furusho M, Miller RH, Bansal R (2012) ERK1/ERK2 MAPK signaling is required to increase myelin thickness independent of oligodendrocyte differentiation and initiation of myelination. J Neurosci 32:8855–8864.

Jeffries MA, Urbanek K, Torres L, Wendell SG, Rubio ME, Fyffe-Maricich SL (2016) ERK1/2 Activation in Preexisting Oligodendrocytes of Adult Mice Drives New Myelin Synthesis and Enhanced CNS Function. J Neurosci 36:9186–9200.

Johansen-Berg H, Baptista CS, Thomas AG (2012) Human structural plasticity at record speed. Neuron 73:1058–1060.

Knott GW, Quairiaux C, Genoud C, Welker E (2002) Formation of dendritic spines with GABAergic synapses induced by whisker stimulation in adult mice. Neuron 34:265–273.

Kuhlman SJ, O’Connor DH, Fox K, Svoboda K (2014) Structural plasticity within the barrel cortex during initial phases of whisker-dependent learning. J Neurosci 34:6078–6083.

Lonze BE, Ginty DD (2002) Function and regulation of CREB family transcription factors in the nervous system. Neuron 35:605–623.

Mayr B, Montminy M (2001) Transcriptional regulation by the phosphorylation-dependent factor CREB. Nat Rev Mol Cell Biol 2:599–609.

McKenzie IA, Ohayon D, Li H, de Faria JP, Emery B, Tohyama K, Richardson WD (2014) Motor skill learning requires active central myelination. Science 346:318–322.

Meffre D, Massaad C, Grenier J (2015) Lithium chloride stimulates PLP and MBP expression in oligodendrocytes via Wnt/beta-catenin and Akt/CREB pathways. Neuroscience 284:962–971.

Mensch S, Baraban M, Almeida R, Czopka T, Ausborn J, El Manira A, Lyons DA (2015) Synaptic vesicle release regulates myelin sheath number of individual oligodendrocytes in vivo. Nat Neurosci 18:628–630.

Michel K, Zhao T, Karl M, Lewis K, Fyffe-Maricich SL (2015) Translational control of myelin basic protein expression by ERK2 MAP kinase regulates timely remyelination in the adult brain. J Neurosci 35:7850–7865.

Norrmen C, Suter U (2013) Akt/mTOR signalling in myelination. Biochem Soc Trans 41:944–950.

Paxinos G, Watson C (1998) The rat brain in stereotaxic coordinates, 4th Edition. San Diego: Academic Press.

Piscopo DM, Weible AP, Rothbart MK, Posner MI, Niell CM (2018) Changes in white matter in mice resulting from low-frequency brain stimulation. Proc Natl Acad Sci U S A 115:E6339–E6346.

Rocamora N, Welker E, Pascual M, Soriano E (1996) Upregulation of BDNF mRNA expression in the barrel cortex of adult mice after sensory stimulation. J Neurosci 16:4411–4419.

Sagi Y, Tavor I, Hofstetter S, Tzur-Moryosef S, Blumenfeld-Katzir T, Assaf Y (2012) Learning in the fast lane: new insights into neuroplasticity. Neuron 73:1195–1203.

Sampaio-Baptista C, Johansen-Berg H (2017) White Matter Plasticity in the Adult Brain. Neuron 96:1239–1251.

Sampaio-Baptista C, Khrapitchev AA, Foxley S, Schlagheck T, Scholz J, Jbabdi S, DeLuca GC, Miller KL, Taylor A, Thomas N, Kleim J, Sibson NR, Bannerman D, Johansen-Berg H (2013) Motor Skill Learning Induces Changes in White Matter Microstructure and Myelination. The Journal of Neuroscience 33:19499–19503.

Scholz J, Klein MC, Behrens TE, Johansen-Berg H (2009) Training induces changes in white-matter architecture. Nat Neurosci 12:1370–1371.

Sinclair JL, Fischl MJ, Alexandrova O, Hebeta M, Grothe B, Leibold C, Kopp-Scheinpflug C (2017) Sound-Evoked Activity Influences Myelination of Brainstem Axons in the Trapezoid Body. J Neurosci 37:8239–8255.

Smith SM, Jenkinson M, Johansen-Berg H, Rueckert D, Nichols TE, Mackay CE, Watkins KE, Ciccarelli O, Cader MZ, Matthews PM, Behrens TE (2006) Tract-based spatial statistics: voxelwise analysis of multi-subject diffusion data. Neuroimage 31:1487–1505.

Smith SM, Jenkinson M, Woolrich MW, Beckmann CF, Behrens TE, Johansen-Berg H, Bannister PR, De Luca M, Drobnjak I, Flitney DE, Niazy RK, Saunders J, Vickers J, Zhang Y, De Stefano N, Brady JM, Matthews PM (2004) Advances in functional and structural MR image analysis and implementation as FSL. Neuroimage 23 Suppl 1:S208–219.

Snaidero N, Mobius W, Czopka T, Hekking LH, Mathisen C, Verkleij D, Goebbels S, Edgar J, Merkler D, Lyons DA, Nave KA, Simons M (2014) Myelin membrane wrapping of CNS axons by PI(3,4,5)P3-dependent polarized growth at the inner tongue. Cell 156:277–290.

Trachtenberg JT, Chen BE, Knott GW, Feng G, Sanes JR, Welker E, Svoboda K (2002) Long-term in vivo imaging of experience-dependent synaptic plasticity in adult cortex. Nature 420:788–794.

Tseng KC, Li H, Clark A, Sundem L, Zuscik M, Noble M, Elfar J (2016) 4-Aminopyridine promotes functional recovery and remyelination in acute peripheral nerve injury. EMBO Mol Med 8:1409–1420.

Vandesompele J, De Preter K, Pattyn F, Poppe B, Van Roy N, De Paepe A, Speleman F (2002) Accurate normalization of real-time quantitative RT-PCR data by geometric averaging of multiple internal control genes. Genome Biol 3:RESEARCH0034.

von Heimendahl M, Itskov PM, Arabzadeh E, Diamond ME (2007) Neuronal activity in rat barrel cortex underlying texture discrimination. PLoS Biol 5:e305.

Williams C, Mehrian Shai R, Wu Y, Hsu YH, Sitzer T, Spann B, McCleary C, Mo Y, Miller CA (2009) Transcriptome analysis of synaptoneurosomes identifies neuroplasticity genes overexpressed in incipient Alzheimer’s disease. PLoS One 4:e4936.

Winkler AM, Ridgway GR, Webster MA, Smith SM, Nichols TE (2014) Permutation inference for the general linear model. Neuroimage 92:381–397.

Winkler AM, Webster MA, Brooks JC, Tracey I, Smith SM, Nichols TE (2016) Non-parametric combination and related permutation tests for neuroimaging. Hum Brain Mapp 37:1486–1511.

Zhang W, Liu HT (2002) MAPK signal pathways in the regulation of cell proliferation in mammalian cells. Cell Res 12:9–18.

