## Supplementary Fig. 1 for "Experience alterations in white matter structure and myelin-related gene expression in adult rats"

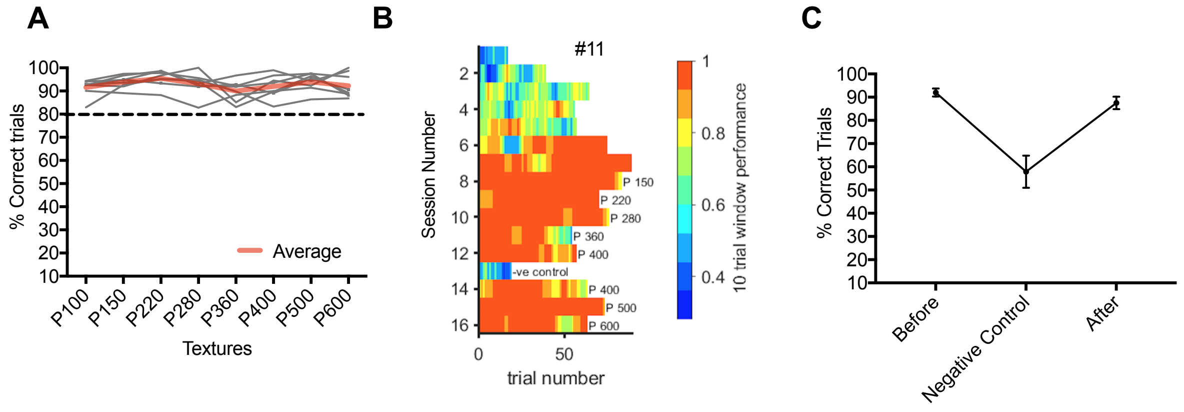


**Supplementary Fig. 1** ***Texture Detection Task (TDT) Performance***. ***A*** *A subgroup of rats (n = 8) were further trained to detect increasingly more fine-grained textures. Individual performance accuracy (% correct trials) for the subset of TDT animals given continued texture detection exposure (n=8; P100 - 162 μm average particle diameter; P150 - 100 μm; P220 - 68 μm; P280 - 52.2 μm; P360 - 40.5 μm; P400 - 35 μm; P500 - 30.2 μm and P600 - 25.8 μm).* After the rats had associated the correct reward side with the first texture, increasing the difficulty of the texture discrimination did not alter their accuracy. *Red line represents average group accuracy.* ***B*** *Graph represents sessions of a rat trained until the smoothest texture (P600). Performance improves over early days and remains high as task difficulty increases. Colour-coding performance scale shown on the right.* ***C*** *Negative control experiment was performed on session 13. When animals were presented with the same texture (P400 versus P400) their performance accuracy dropped to chance levels. RM- Anova $F_{(2,14)} = 16.897$, $p < 0.0001$. Planned comparison paired T-Test between performance before and NC $t_{(7)} = 4.883$, $p < 0.01$; performance after and NC $t_{(7)} = -4.163$, $p < 0.01$). Before – Performance in the TDT task before Negative Control, After – Performance in the TDT task after the Negative Control.*
